# Oxaliplatin kills cells via liquid-liquid demixing of nucleoli

**DOI:** 10.1101/2021.06.10.447918

**Authors:** H. Broder Schmidt, Zane A. Jaafar, Jason J. Rodencal, Manuel D. Leonetti, Scott J. Dixon, Rajat Rohatgi, Onn Brandman

## Abstract

Platinum (Pt) compounds such as oxaliplatin are amongst the most commonly prescribed anti-cancer drugs. Despite their considerable clinical impact, the molecular basis of platinum cytotoxicity and cancer specificity remain unclear. Here, we show that oxaliplatin, a backbone for the treatment of colorectal cancer, causes liquid-liquid demixing of nucleoli at clinically-relevant concentrations by interfering with the interaction networks that organize nucleoli. This biophysical defect leads to cell cycle arrest, impaired rRNA processing and shutdown of PolI-mediated transcription, ultimately resulting in cell death. We propose that the mechanism of action of oxaliplatin provides a blueprint for the therapeutic targeting of the increasing number of cellular processes being linked to biomolecular condensates.

## MAIN TEXT

Platinum (Pt) compounds (Fig. S1) were discovered serendipitously over fifty years ago when the electrolysis products of platinum electrodes were shown to inhibit the growth of *E. coli* (*1*). Cisplatin, the first platinum compound to be developed as a result of these pioneering studies, soon became the cornerstone of treatment for ovarian, lung, head and neck, bladder and germ cell cancers (*2*). Significant toxicities, including kidney damage, nausea and vomiting, hearing changes and peripheral neuropathy, led to the development of additional platinum analogs, two of which (carboplatin and oxaliplatin) are used clinically (*1*). Oxaliplatin is particularly effective in colorectal cancer, a disease in which cisplatin and carboplatin have no meaningful activity (*2*). Understanding the mechanism of action (MOA) of oxaliplatin and other platinum compounds promises to unlock new principles in how these highly reactive (and relatively non-specific) agents kill specific cancer cells with a clinically acceptable therapeutic window.

While the highly reactive platinum warhead common to Pt drugs can form adducts with all classes of cellular macromolecules (DNA, RNA and proteins) (*3*), the dominant view is that their cytotoxicity is caused by the formation of intrastrand adducts with purine bases in DNA, ultimately leading to failed DNA-damage responses (*4*). This model is partly based on the correlation between cytotoxicity and the abundance of specific G-G and A-G DNA-Pt adducts. In contrast, recent biochemical and drug profiling studies categorized oxaliplatin, but not cisplatin, as a transcription-translation inhibitor and ascribed its cytotoxic effects to ribosome biogenesis stress (*5, 6*). However, the MOA by which oxaliplatin inhibits ribosome biogenesis remains elusive.

To explore how oxaliplatin affects ribosome function in primary cells, we performed Ribo-seq, which measures total and ribosome-protected RNA, in unperturbed and oxaliplatin-treated human umbilical vein endothelial cells (HUVECs) (*7*). This allowed us to detect changes in both transcript abundance and translation efficiency at single nucleotide resolution. Surprisingly, oxaliplatin treatment increased the translation efficiency of transcripts that encode ribosomal proteins (Fig. 1A; Supplementary File 1) and the abundance of transcripts encoding small nucleolar RNAs (snoRNAs), in particular *SNORD* and *SNORA* (Fig. 1B; Supplementary File 1), suggesting that oxaliplatin induced nucleolar stress (*8*). Indeed, GO term analysis revealed that the top cellular component associated with the transcripts upregulated in oxaliplatin-treated HUVECs was the nucleolus (Fig. 1B).

**Figure 1.**
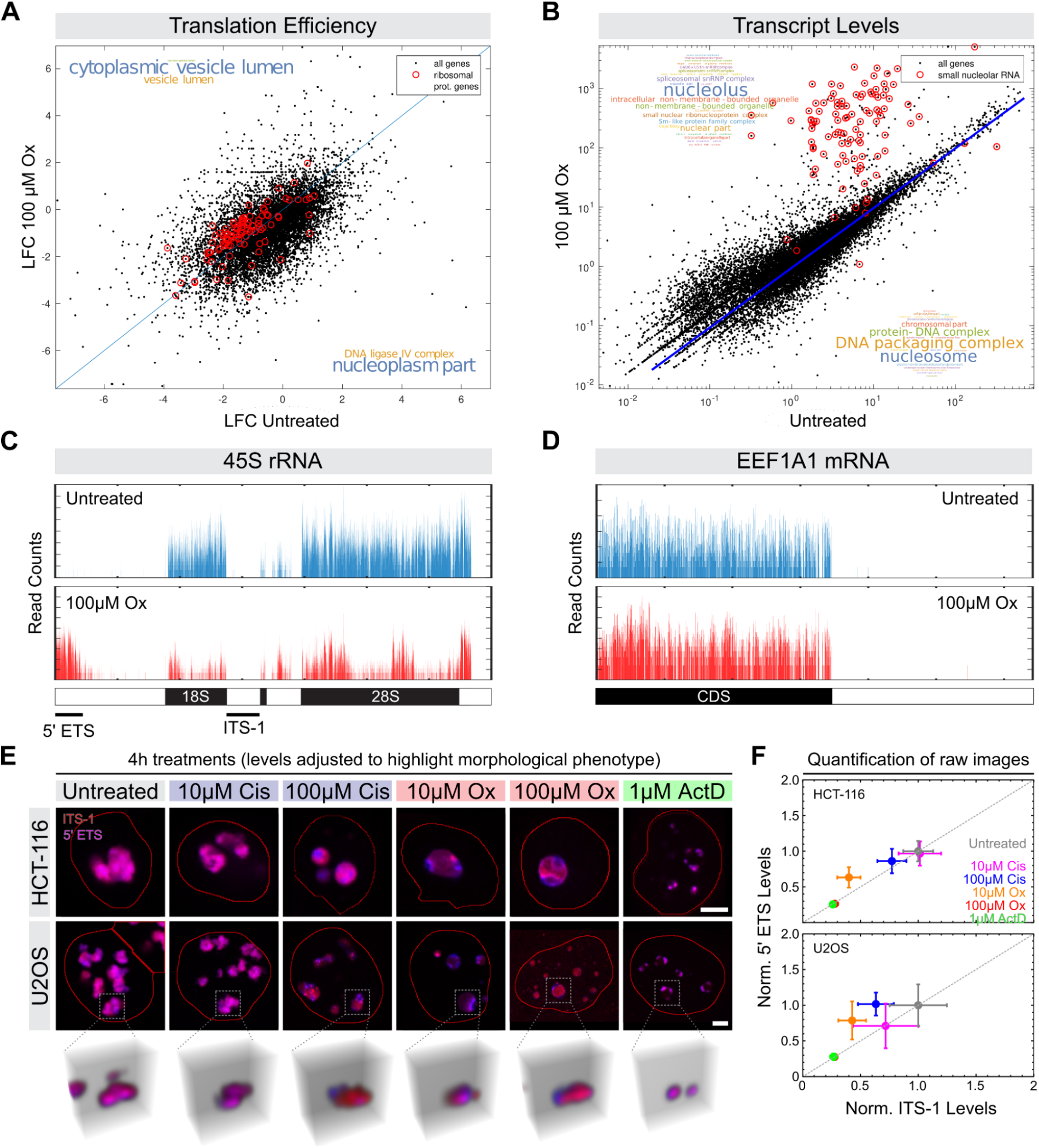
Oxaliplatin stabilizes pre-rRNA transcripts and alters sub-nucleolar rRNA localization. (**A**) mRNA translation efficiency, calculated as log_2_(median normalized ribosome footprint frequency/mRNA frequency), in HUVECs treated with and without 100 μM oxaliplatin for 4 hours. Inserts show word clouds for component GO terms associated with mRNAs that are translated more (top) or less (bottom) efficiently in oxaliplatin-treated HUVECs. (**B**) RNA transcript levels in HUVECs treated with and without 100 μM oxaliplatin for 4 hours. Inserts show word clouds for component GO terms associated with transcripts that are up- (top) or down- (bottom) regulated in oxaliplatin-treated HUVECs. (**C**) and (**D**) Mapping of RNA reads to the 45S rRNA (C) and *EEF1A1* mRNA (D) in untreated (blue) and oxaliplatin-treated (red) HUVECs. Cartoons below outline transcript architecture. The black bars in (C) indicate the 5’ ETS and ITS-1 regions targeted by FISH. (**E**) Overlays of representative confocal images showing RNA-FISH stainings against the 5’ ETS (blue) and ITS-1 (red) in HCT-116 and U2OS cells treated with the indicated amounts of cisplatin (Cis), Oxaliplatin (Ox) and actinomyosin-D (ActD) for 4 hours. Image levels are individually adjusted to highlight morphological phenotypes. See Figure S4 for raw data. The bottom row shows 3D reconstructions of the indicated nucleoli in U2OS cells based on confocal z-stacks. Scale bar: 5 μm. (**F**) Quantification of the nucleolar 5’ ETS (y-axis) and ITS-1 (x-axis) levels in raw images of treated and untreated HCT-116 and U2OS cells (N ≥ 75 nucleoli per condition). Plot markers indicate mean values and error bars the standard deviation.

Nucleoli are phase-separated, multi-layered protein and RNA condensates with liquid-like properties (Fig. S2) that are required for rRNA transcription, rRNA processing, and ribosome biogenesis (*9*). This prompted us to analyze the effect of oxaliplatin on rRNA transcripts in more detail. The majority of rRNAs are derived from a large, noncoding precursor called the 45S rRNA, which is first synthesized by RNA polymerase I (PolI) and then cleaved into mature 28S, 18S and 5.8S rRNA molecules, two steps that take place in nucleoli (*10*). When we mapped transcript reads to the 45S rRNA, we observed that they localized almost exclusively to the 28S, 18S and 5.8S rRNA regions in untreated cells (Fig. 1C). We also noticed additional, unexpected reads corresponding to the 5’ external transcribed sequence (5’ ETS) region only in oxaliplatin-treated cells (Fig. 1C). The 5’ ETS is a short-lived intermediate in 45S rRNA processing that normally does not accumulate in the cytosol. This effect of oxaliplatin was specific to the 45S since we did not observe additional signals in protein-coding transcripts like EEF1A1 (Fig. 1D). Our ability to detect reads mapping to the 5’ ETS depended on the use of library circularization, a routine tep in the Ribo-seq protocol (Fig. S3). Since oxaliplatin-RNA adducts can block the library amplification step of standard RNA-seq protocols, circularization enables library generation from RNA fragments regardless of whether they contain platinum adducts (*11*). This indirectly suggests that RNA fragments mapping to the 5’ ETS may contain oxaliplatin adducts. Our results show that oxaliplatin disrupts the processing of rRNA from its 45S precursor and stabilizes normally short-lived intermediates like the 5’ ETS.

To corroborate this finding, we performed fluorescence in situ hybridization (FISH) against the 5’ ETS and the downstream internal transcribed sequence-1 (ITS-1) that separates the 18S and 5.8S rRNAs (Fig. 1C). As expected (*12*), both the 5’ ETS and spacer signals co-localized and showed a characteristic nucleolar staining in untreated human colon cancer (HCT-116) and osteosarcoma (U2OS) cell lines (Figs. 1E; S4). By contrast, in cells treated with high doses of oxaliplatin and cisplatin (100 μM), the 5’ ETS and ITS-1 segregated into distinct regions of highly spherical nucleoli, with the 5’ ETS confined to the periphery and sharing little overlap with the ITS-1 (Figs. 1E; S4). Notably, oxaliplatin induced these phenotypes at 10-fold lower concentrations across all cell lines tested compared to cisplatin (Fig. 1E). Low doses of the RNA polymerase inhibitor actinomycin D (1 μM), a nucleolar-stress inducing agent that interferes with rRNA transcription but not processing (*12*), also changed the sub-nucleolar distribution of the 5’ ETS and ITS-1, but did not cause their segregation, as observed with oxaliplatin (Fig. 1E). Quantification of the FISH signals confirmed that oxaliplatin treatment stabilized the 5’ ETS, despite a decline in overall levels of the 45S rRNA (Fig. 1F). We conclude that oxaliplatin disrupts the core functions of nucleoli, in particular PolI-mediated transcription and 45S rRNA processing.

Given the oxaliplatin-induced interference with nucleolar function that we observed and the recent finding that certain antineoplastic drugs readily partition into nuclear condensates such as nucleoli (*13*), we sought to test how oxaliplatin affects nucleolar ultrastructure. To this end, we endogenously tagged the nucleolar scaffold components FBL and NPM1 with fluorescent markers in U2OS cells. FBL is a major constituent of the dense fibrillar center (DFC), the sub-nucleolar site of rRNA processing, while NPM1 is a marker of the granular component (GC) where ribosomal subunits are assembled (*14*). When treated with oxaliplatin, we observed an increase in nucleolar sphericity and a dramatic sub-nucleolar segregation of major nucleolar proteins (FBL, NPM1) and rRNA (5’ ETS) (Figs. 2A-C; S5A). Notably, the 5’ ETS was enriched at the interface of two nearly completely separated FBL and NPM1 phases (Fig. 2A, B), suggesting a dramatic change in nucleolar surface tension. Indeed, by measuring nucleolar surface fluctuations (*15*), we estimated that the surface tension at the NPM1-nucleoplasm interface increased by an order of magnitude in oxaliplatin-treated cells (Fig. 2D), demonstrating that oxaliplatin interferes with the multiphase behavior of nucleoli.

**Figure 2.**
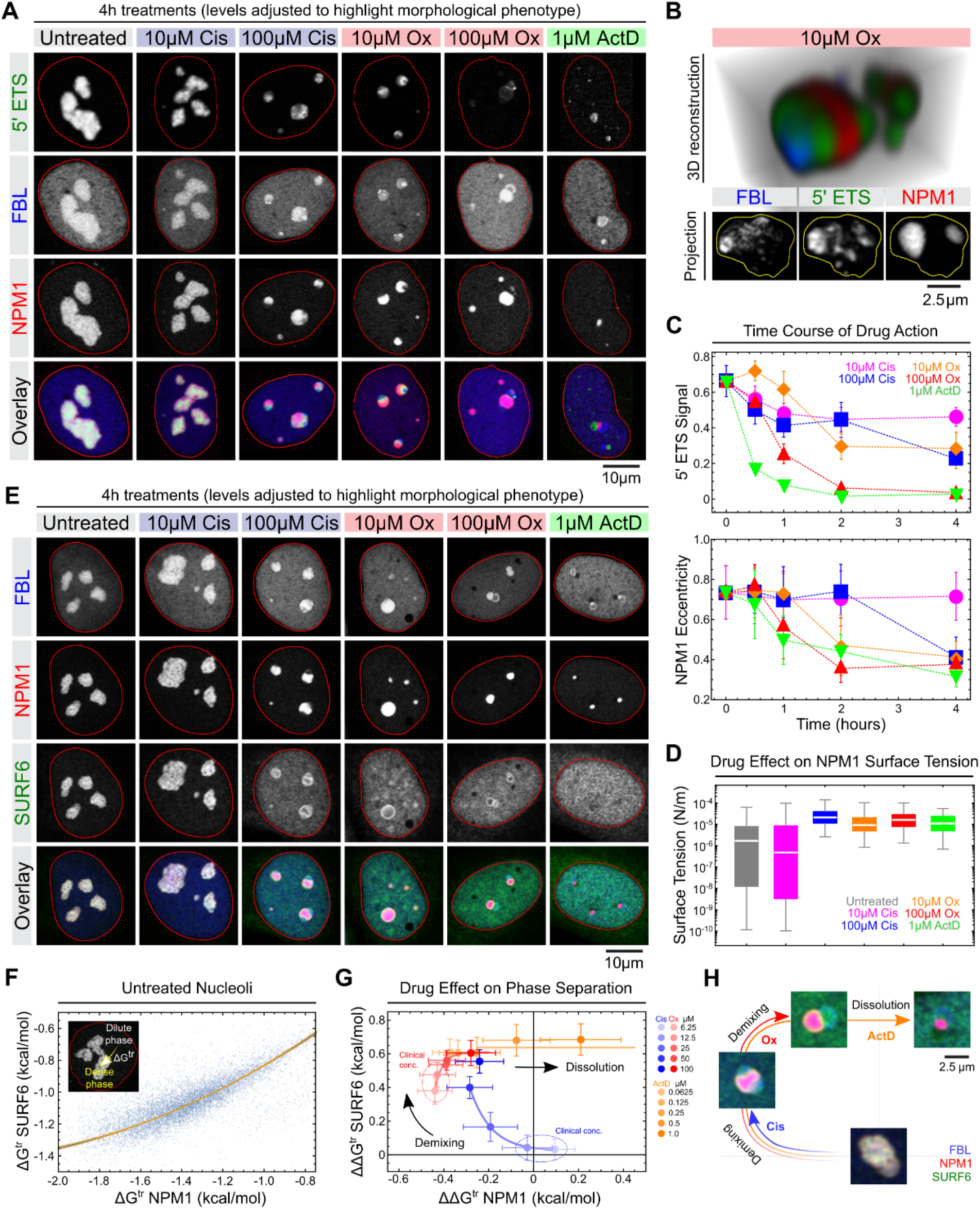
Oxaliplatin causes the disintegration of nucleolar substructure. (**A**) Representative confocal images of RNA-FISH stainings against the 5’ ETS (green) in U2OS-Rainbow cells expressing FBL-Halo (blue) and NPM1-RFP (red) from their endogenous loci. The cells were treated with the indicated amounts of drugs for 4 hours prior to staining. Image levels were individually adjusted to highlight morphological phenotypes (see Figure S5 for unmodified images). (**B**) Multi-channel 3D reconstruction and 2D projection of a representative nucleolus in cells treated with 10 μM oxaliplatin (Ox) for 4 hours. Nuclear background signals were filtered out to enhance contrast (see Methods). (**C**) Quantification of the decline of 5’ ETS signal and NPM1 eccentricity in U2OS-Rainbow cells treated with the indicated amounts of drugs over time. Plot markers indicate mean values and error bars the standard deviation. At least 106 nucleoli per condition and time point were quantified in raw images. (**D**) Quantification of the changes in NPM1 surface tension in response to drug treatments. Horizontal lines depict median values, boxes the 25%-75% and error bars the 5%-95% percentiles. N = 136 (untreated), 200 (10 μM Cis), 217 (100 μM Cis), 178 (10 μM Ox), 167 (100 μM Ox) and 40 (1 μM ActD). (**E**) Representative confocal images of immunostaining against SURF6 (green) in U2OS-Rainbow cells expressing FBL-Halo (blue) and NPM1-RFP (red) after 4 hour treatments with the indicated amount of drugs. Image levels were individually adjusted to highlight morphological phenotypes (see Figure S5 for unmodified images). (**F**) Dependence of the NPM1 and SURF6 transfer energies from the dilute phase (nucleoplasm) into the dense phase (nucleoli) in untreated UCSF-Rainbow cells immunostained for SURF6 (N = 10,893). (**G**) Change of NPM1 and SURF6 transfer energies upon treatment with the indicated amounts of cisplatin (blue shades), oxaliplatin (red shades) and actinomycin D (orange shades) for 4 hours relative to untreated cells. The dashed circles indicate clinical concentrations of cisplatin and oxaliplatin. At least 2,132 nucleoli per condition were quantified. (**H**) Schematic summarizing panels E and G to outline how cisplatin, oxaliplatin and actinomycin D affect nucleolar morphology.

Whereas nucleolar surface tension increased in response to both platinum compounds, oxaliplatin caused the demixing of the nucleolar phases at ten-fold lower concentrations and four-fold earlier time points compared to cisplatin (Figs. 2A, C). Unique to oxaliplatin, at higher doses (100 uM) the segregation of the nucleolar layers was so pronounced that FBL formed characteristic ‘nucleolar caps’ (Figs. 2A; S5A) associated with transcriptional arrest and cellular stress (*16, 17*). In comparison to the Pt drugs, actinomycin D caused an almost complete separation of the 5’ ETS and protein markers (Fig. 2A). Time course analysis revealed that actinomycin D first decreased 5’ ETS levels and then altered nucleolar morphology, whereas oxaliplatin affected both in concert (Fig. 2C). This observation is consistent with the model that oxaliplatin alters nucleolar morphology, thereby causing a secondary effect on rRNA transcription. Together, our data show that oxaliplatin causes the demixing of the DFC and GC phases at clinically relevant concentrations (~5-10 μM, (*5*)) via a mechanism that is distinct from transcriptional inhibition by actinomycin D.

The morphology and multiphase organization of nucleoli is the result of an elaborate network of heterotypic interactions between multiple nucleolar proteins and rRNA (*18, 19*). Consequently, disruption of this interaction network may underlie the nucleolar demixing seen in oxaliplatin-treated cells. To test this model, we focused on the well-studied interaction between NPM1 and the non-ribosomal GC component SURF6, which is important for both the nucleolar localization and phase separation behavior of NPM1 (*20, 21*). While NPM1 and SURF6 co-localize in untreated cells, oxaliplatin treatment both greatly reduced the nucleolar localization of SURF6 and caused a nearly complete demixing of NPM1 and the remaining nucleolar SURF6 (Figs. 2D, E; S5B). Strikingly, SURF6 accumulated at the surface of the NPM1- and FBL-rich phase or within the nucleolar caps (Fig. 2D), again reflecting dramatic changes in surface tension in response to oxaliplatin treatment. To gain quantitative insights into the oxaliplatin-induced changes in the NPM1-SURF6 interaction and phase separation, we measured the transfer energies ΔG^tr^ of NPM1 and SURF6 from the nucleoplasm (dilute phase) into nucleoli (dense phase), as previously described (*19, 20*). We observed that ΔG^tr^ of NPM1 and SURF6 are strongly dependent in untreated cells (Fig. 2F), as expected for two proteins that phase separate together (*19–21*). Using ΔG^tr^ in untreated cells as a reference, we then determined the changes in ΔG^tr^ induced by increasing concentrations of oxaliplatin, cisplatin and actinomycin D. While clinical doses of cisplatin (<12.5 μM; (*5*)) hardly affected ΔG^tr^ of neither NPM1 nor SURF6, concentrations greater than 25 μM simultaneously caused ΔG^tr^ (NPM1) to become more negative and ΔG^tr^ (SURF6) less negative (Fig. 2G). This is consistent with the destabilization of the NPM1-SURF6 interaction and demixing of the GC phase due to increased NPM1 and decreased SURF6 phase separation (Fig. 2H). Notably, oxaliplatin treatment caused these effects at concentrations as low as 6.25 μM (Fig. 2G). Above concentrations of 50 μM, oxaliplatin started to reduce the driving force for NPM1 phase separation again. However, only treatment with increasing doses of actinomycin D caused the values of both ΔΔG^tr^ (NPM1) and ΔΔG^tr^ (SURF6) to become positive, indicative of nucleolar dissolution (Fig. 2G, H). We conclude that oxaliplatin causes the demixing of individual nucleolar phases at clinical concentrations and suggest that this is due to the disruption of the heterotypic interactions between proteins and rRNA that constitute these phases.

We next sought to determine how oxaliplatin interferes with nucleolar interaction networks. Recent work showed that the formation and integrity of multiphase protein-RNA condensates such as nucleoli and stress granules are dictated by the valency and stoichiometry of key nodes in the condensate interaction networks (*19, 22–24*). Thus, such networks can be disrupted by removing central nodes or changing their valency. Pt compounds are known to modify and crosslink nucleic acids and proteins (*25*), suggesting that the chemical modification of nucleolar components with platinum may directly interfere with their valency. To test this model, we purified recombinant FBL and NPM1 from bacteria and incubated the proteins with varying concentrations of oxaliplatin and cisplatin. We found that oxaliplatin readily cross-linked FBL and, to a lesser extent, NPM1 in a dose-dependent manner, whereas cisplatin had little effect (Fig. 3A and B). When we mixed FBL with rRNA to form condensates that mimic the DFC, oxaliplatin preferentially platinated FBL over rRNA (Fig. 3C). In contrast, we observed that oxaliplatin had little effect on NPM1-rRNA condensates that mimic the GC, whereas cisplatin introduced rRNA crosslinks under these conditions (Fig. 3D). We confirmed that phase separation enhanced the respective protein- and nucleic acid-modifying activities of oxaliplatin and cisplatin by forming condensates of FBL and NPM1 with various nucleic acids (Fig. S6). These findings suggest that oxaliplatin preferentially targets and modifies FBL, thereby triggering a cascade that disrupts the multiphase organization of nucleoli at large.

**Figure 3.**
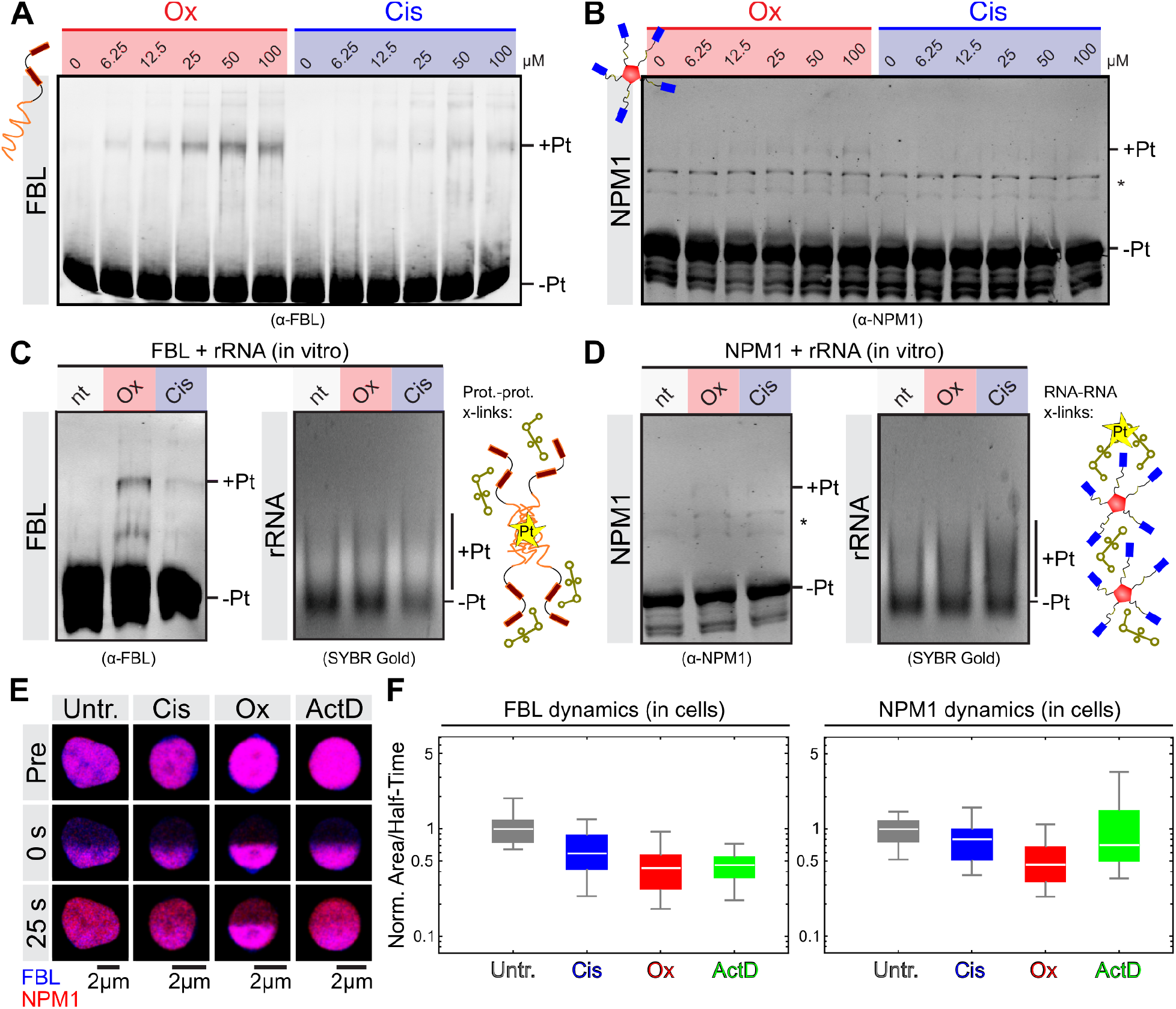
Oxaliplatin preferentially modifies the dense fibrillar component marker FBL. (**A**) and (**B**) Gel shifts assays to detect the dose-dependent cross-linking of recombinant FBL (A) and NPM1 (B) by oxaliplatin (Ox) and cisplatin (Cis) visualized by immunoblotting. Labels mark the major bands corresponding to unmodified (-Pt) and platinated (+Pt) FBL and NPM1. Asterisks denote unspecific bands. Reactions were incubated for 20 hours at room temperature. (**C**) and (**D**) Gel shifts assays to detect protein and rRNA cross-linking in FBL-rRNA (A) and NPM1-rRNA (B) mixtures after treatment for 20 hours with 100 μM oxaliplatin or cisplatin. Proteins were visualized by immunoblotting and rRNA by SYBR gold staining. Labels as in (A) and (B). (**E**) Half-bleach experiments in live U2OS-Rainbow cells to compare nucleolar dynamics. Representative overlays of FBL (blue) and NPM1 (red) channels before bleaching (pre), immediately after (0 s) and during recovery (25 s). Image levels were adjusted and nuclear background was removed to enhance contrast (see Figure S7 for unmodified images). (**F**) Quantification of nucleolar half-bleach experiments. Plots show the diffusion rate (bleached area over recovery half-time) of FBL and NPM1 relative to untreated nucleoli. Horizontal lines depict median values, boxes the 25%-75% percentiles and error bars the min and max values. N = 19 (ActD), 27 (Cis), 23 (Ox), 25 (Untreated).

The striking morphological changes in nucleolar substructure and small extent of cross-linking in vitro suggest that oxaliplatin introduces post-translational modifications (PTMs) that maintain (but may alter) the overall liquid nature of nucleoli, rather than stable intermolecular bonds that abolish nucleolar dynamics. To test this, we simultaneously photobleached the FBL and NPM1 signal in one half of control or treated nucleoli and measured fluorescence recovery over time (Figs. 3E; S7). Given that our drug treatments affect the size of nucleoli, we determined the diffusion coefficients (bleached area divided by the half-time of fluorescence recovery) of FBL and NPM1 to quantitatively compare the data. While oxaliplatin treatment reduced both FBL and NPM1 dynamics by more than half, cisplatin and actinomycin D only affected FBL dynamics (Fig. 3F). This further demonstrates that oxaliplatin and actinomycin D act via different mechanisms. We propose that oxaliplatin interferes with the nucleolar interaction network by direct chemical modification of nucleolar proteins and rRNA, whereas the transcription inhibitor actinomycin D does so by removing rRNA, a key node of the interaction network (*19*).

In addition to the scaffold components that drive phase separation and define material properties, biomolecular condensates like nucleoli also contain various client molecules that interact with the scaffolds (*26*). To test how oxaliplatin affects client recruitment to nucleoli, we looked at the cell proliferation and cancer prognosis marker KI67 (*27*). In untreated cells, KI67 localized to the nucleolar rim (Figs. 4A; S8), as previously reported (*28*). Given the surfactant-like properties of KI67 (*29*), we expected that its association with nucleoli would be highly sensitive to the changes in nucleolar surface tension induced by oxaliplatin. Indeed, we found that low doses of oxaliplatin caused the loss of KI67 from the nucleolar rim and its dispersion into the nucleoplasm (Fig. 4B, C), providing further evidence that platination alters both biochemical interactions and biophysical forces within nucleoli.

**Figure 4.**
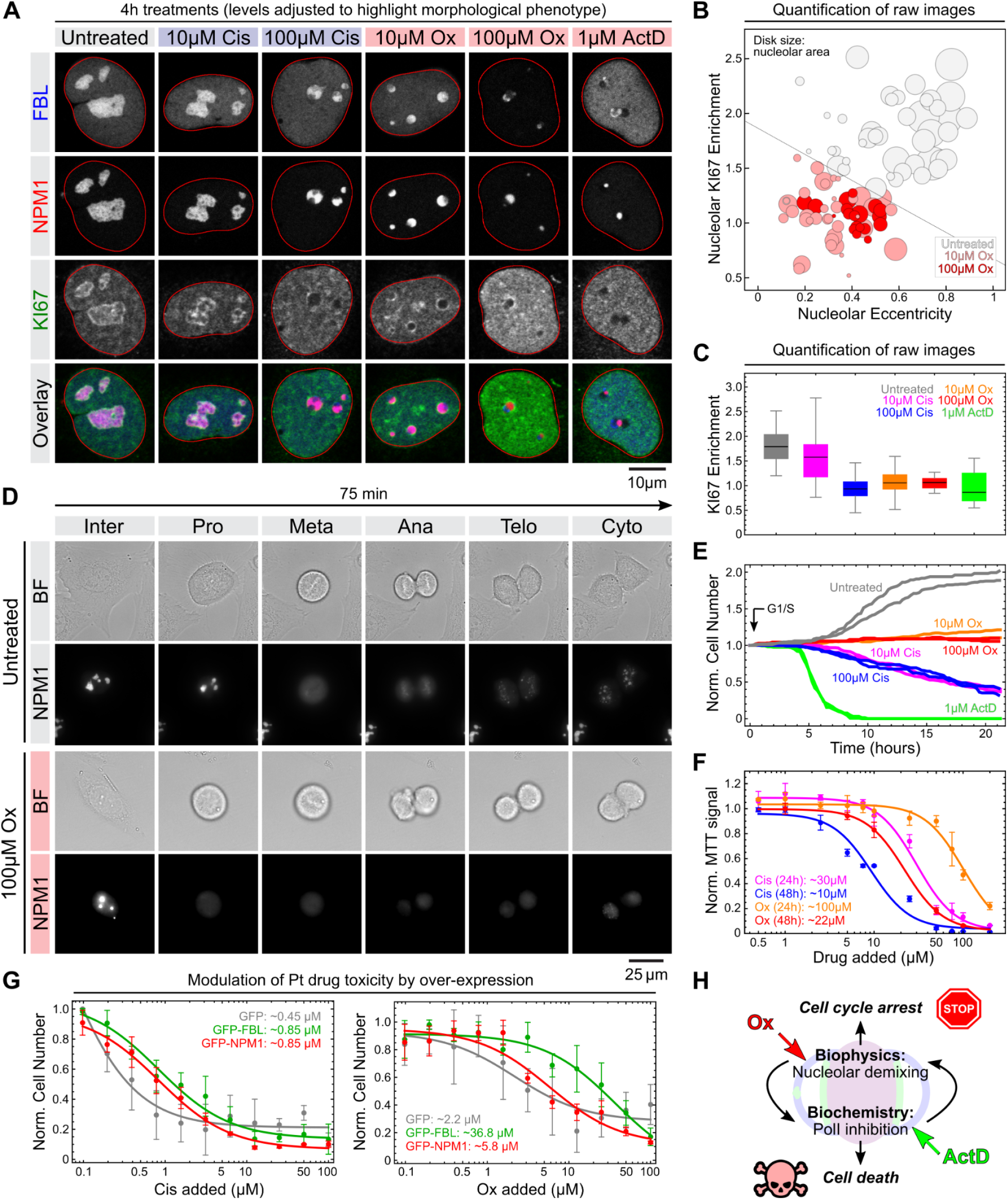
Oxaliplatin treatment results in the loss of nucleolar clients, cell cycle defects and cell death. (**A**) Representative confocal images of U2OS-Rainbow cells immunostained for the nucleolar rim and cell proliferation marker KI67 (green) after 4 hour treatments with the indicated amount of drugs. Image levels were individually adjusted to highlight morphological phenotypes (see Figure S8 for unmodified images). (**B**) Quantification of nucleolar KI67 enrichment (y-axis), nucleolar eccentricity (x-axis) and nucleolar area (plot marker size) in oxaliplatin-treated cells. At least 20 nucleoli per condition were analyzed. (**C**) Quantification of nucleolar KI67 enrichment in oxaliplatin-, cisplatin- and actinomycin D-treated cells. At least 15 nucleoli per condition were analyzed. (**D**) Live cell imaging of nucleolar dynamics during mitosis in control and oxaliplatin-treated U2OS cells. (**E**) Quantification of cell number during live cell imaging of control and drug-treated cells for 24 hours. At least 73 cells per replicate were tracked over time. (**F**) MTT assay to measure metabolic activity of cells treated with the indicated amounts of Pt compounds for 24 and 48 hours. Data points represent the mean values from four replicates, error bars the standard deviation and curves a nonlinear dose-response fit to the data (see Methods). Numbers indicate the EC50 values derived from the fits. (**G**) Cell number of HT-1080 cells transfected with either GFP, GFP-FBL or GFP-NPM1 after 48 hour treatments with the indicated amounts of Pt compounds. Data is plotted as in (F). Numbers indicate the EC50 values. (**H**) Illustration of the different mechanisms by which oxaliplatin and actinomycin D target nucleoli and cause cell death.

During mitosis, nucleoli are disassembled and KI67 is localized to mitotic chromosomes, where it plays a key role in chromosome dispersal and nuclear reassembly (*29, 30*). Notably, KI67 chromosome association is also a prerequisite for NPM1 recruitment and the reformation of nucleoli (*31*). This prompted us to investigate the effect of oxaliplatin on nucleoli during mitosis in synchronized U2OS cells with live cell imaging (Fig. S9). Using endogenously-tagged NPM1 as a marker, we found that the nucleoli of both untreated and oxaliplatin-treated cells readily disassemble following nuclear envelope breakdown at the end of prophase (Fig. 4D). However, in oxaliplatin-treated cells, NPM1 failed to localize to mitotic chromosomes during anaphase and nucleoli failed to reform during telophase (Fig. 4D). In agreement with previous reports (*32*), only few oxaliplatin-treated cells entered the cell cycle (Fig. 4E). Prolonged oxaliplatin treatment eventually caused cell death, as measured by metabolic activity (Fig. 4F). We could not observe nucleolar reformation in cisplatin- and actinomycin D-treated cells due to much greater toxicity, including cisplatin concentrations that did not affect nucleoli (Fig. 4E, F).

Given our observation that FBL is readily modified by oxaliplatin, we next explored how FBL overexpression affects cell growth in oxaliplatin-treated cells. To this end, we measured the toxicity of oxaliplatin and cisplatin in HT-1080 cells transfected with GFP-FBL, GFP-NPM1 or GFP alone. FBL overexpression conferred a dramatic resistance to oxaliplatin, resulting in a >15-fold increase in EC50 (Fig. 4G). This effect was specific to oxaliplatin and FBL, as it was not recapitulated with cisplatin or overexpression of NPM1. Our findings show that oxaliplatin causes cell cycle arrest and ultimately cell death, both of which can be alleviated by stabilizing the nucleolar interaction network through FBL overexpression.

In conclusion, our study provides a MOA explaining how oxaliplatin acts as a transcription and translation inhibitor. By chemically modifying key nucleolar scaffold components, including FBL, NPM1 and 45S rRNA, oxaliplatin causes the liquid-liquid demixing of nucleoli and triggers cell cycle arrest. In agreement with recent findings (*19, 33*), this biophysical defect then impairs the function of nucleoli to supply cells with rRNA and ribosomes, ultimately leading to cell death (Figs. 4H; S10). We propose that the bulkiness and hydrophobicity of oxaliplatin allow it to preferentially enrich in nucleoli, where it then readily modifies nucleolar RNAs and proteins (*13, 34*). In contrast to small molecule drugs like actinomycin D that specifically target the core transcription machinery, the indirect disruption of PolI-mediated transcription and ribosome biogenesis by nucleolar demixing may expand the therapeutic window of transcription-translation inhibitors. Our findings demonstrate that expanding the arsenal of small molecules targeting biomolecular condensates is a promising strategy for developing the next generation of antineoplastic drugs (*35*).

## ACKNOWLEDGMENTS

We thank Steven Boeynaems, Achuthan Venkatesh, Keren Lasker and Bede Portz for critical feedback and sharing reagents. We are grateful for the support from the National Institutes of Health (GM118082 to RR and GM115968 to OB) and the Stanford Cancer Institute (Innovation Award to OB). JJR was supported by a NIH T32 Cancer Biology Training Grant (5T32CA009302).

## SUPPLEMENTARY MATERIALS

### Materials and Methods

#### Small Molecules and Reagents

**Table S1.** Key commercial reagents used in this study.

| Reagent | Source | Identifier |
| --- | --- | --- |
| Oxaliplatin | Acros Organics | AC456131000 |
| Cisplatin | Biotang | RYG01 |
| Actinomycin-D | MP Biomedicals | 02194525-CF |
| TruSeq Ribo Profile Kit | Illumina | RPYSC12116 |
| 5' ETS FISH probes | Stellaris | Custom* |
| Spacer FISH probes | Stellaris | Custom* |
| Rabbit α-NPM1 polyclonal antibody | Santa Cruz | sc-5564 |
| Rabbit α-FBL monoclonal antibody | Cell Signaling | 2639 |
| Rabbit α-SURF6 polyclonal antibody | Atlas/Sigma | HPA023608 |
| Rabbit α-MKI67 polyclonal antibody | Atlas/Sigma | HPA000451 |
| α-Rabbit-Alexa488 polyclonal antibody | Invitrogen | A-21206 |
| α-Rabbit-Alexa647 polyclonal antibody | Invitrogen | A-31573 |
| α-Rabbit-IR800 polyclonal antibody | LiCor | 926-49020 |

*\*See Table S2 for sequences.*

#### Cell Culture

HCT-116 (ATCC CCL-247) and all U2OS (ATCC HTB-96) cell lines were maintained at 37°C and 5% CO_2_ in DMEM high glucose (GE Healthcare) supplemented with 10% FBS (Atlanta Biologicals), 2mM L-glutamine (Gemini Biosciences), 1mM sodium pyruvate (Gibco), 1x MEM non-essential amino acids solution (Gibco), 40 U/ml penicillin and 40 μg/ml streptomycin (Gemini Biosciences).

HT-1080 (ATCC CCL-121) cells were grown in DMEM high glucose medium (Corning Life Science) supplemented with 1% non-essential amino acids (Life Technologies). HT-1080 cells used in this study were stably infected with Nuc::mKate2, a red fluorescent protein targeted to the nucleus (*36*).

HUVEC were grown and maintained in EGM-2 Endothelial Cell Growth Medium BulletKitTM (Lonza, Catalog Number CC-3162).

#### Generation of U2OS-Rainbow Knock-In Cell Lines

Genome engineering of U2OS cells was performed by co-electroporation of pre-formed Cas9/sgRNA ribonucleo-protein (RNP) complexes and double-stranded homology template donors. Cas9/sgRNA RNPs were assembled in vitro as described (*37*). Homology donors (full sequences below) were obtained by PCR amplification as in (*38*). To enhance homologous recombination, U2OS cells were first synchronized by treatment with 200 ng/mL nocodazole for 20h (*39*). Synchronized cells were harvested from the culture’s supernatant and electroporated in 96-well format using an Amaxa Shuttle nucleofection device (CM-104 program, Lonza). For each individual electroporation, 200,000 cells resuspended in SE solution were mixed with 10 pmole Cas9/sgRNA RNP and 5 pmole homology template. Successfully edited cells were subsequently selected by fluorescence-activated cell sorting (FACS) on a SONY SH800 instrument.

NPM1 donor sequence: https://benchling.com/s/seq-tkR3x4eaRuAhf6FJ5jUU
FBL donor sequence: https://benchling.com/s/seq-n6ABcyxH75cUCoj5aDoc
NPM1 Cas9 gRNA sequence: GCCAGAGATCTTGAATAGCC
FBL Cas9 gRNA sequence: AACTGAAGTTCAGCGCTGTC

#### Ribosome Profiling and RNA-seq

HUVECs were seeded in 15cm plates, grown to near confluence and treated with 100 μM oxaliplatin for 4 hours. The TruSeq Ribo Profile kit (Illumina) was then used to extract ribosome-protected and total, predominantly cytoplasmic RNA from 50 million treated or control cells, and to prepare sequencing libraries for ribosome profiling (Ribo-seq) and expression analysis (RNA-seq) using a circularization step (Fig. S3), respectively. For cytosolic RNA-seq without circularization (Fig. S3), cytosolic RNA fraction was purified according to the protocol in the TruSeq Ribo Profile kit. rRNAs were then depleted using the NEBNext rRNA Depletion Kit (Human/Mouse/Rat) and RNA-seq was performed using the NEBNext Ultra RNA Library Prep Kit for Illumina.

Kallisto (https://pachterlab.github.io/kallisto/) was used to quantify read counts against Refseq GRCh38.p13_rna plus ribosomal rRNA sequences obtained from Ensembl BioMart - (https://www.ensembl.org) for ribosome footprints and total mRNA. Transcripts per million (TPMS) values were then normalized by median values to enable comparison between samples with different total read counts. Values for all genes and conditions are provided in Supplementary File 1.

#### RNA-FISH Staining and Imaging

Cells were seeded onto acid-washed #1.5 glass coverslips (at a density of 50,000 cells/coverslip). After 24 hours, culture medium was exchanged for fresh DMEM containing the indicated amounts of drugs. In case U2OS knock-in lines were used, the cells were stained with 1 μM HaloTag Oregon Green (Promega) in DMEM for 15 minutes at 37°C and washed with PBS prior to drug addition.

Following treatment for 4 hours at 37°C, cells were washed with PBS, fixed with 4% PFA for 15 minutes at room temperature and washed again with PBS. Cells were then incubated in 70% ethanol for 16-24 hours at 4°C, washed with 2x SSC buffer (Ambion) containing 10% formamide (Ambion) and stained with 125 nM dye-labeled FISH probes (Stellaris; see Table S2 for sequences) in 2x SSC buffer (Ambion) containing 10% formamide (Ambion) and 10% (w/v) dextran sulfate (Sigma) for 16 hours at 37°C in a humidity chamber. After two washes with 2x SSC buffer (Ambion) containing 10% formamide (Ambion) for 30 min at 37°C each, cells were mounted onto microscopy slides with ProLong Diamond mounting medium containing DAPI (Molecular Probes). All solutions for RNA-FISH were prepared with nuclease-free water (Ambion).

Stained cells were imaged by taking z-stacks (in 100x steps of 0.45 μm each) using a Leica DMI6000 microscope eeqipped with a Yokogawa CSU-X1 spinning disk head, an Andor iXon Ultra 897 EMCD camera and a 100x (1.4 NA) oil immersion objective. FISH signals were quantified and plotted using custom Mathematica scripts.

**Table S2.**
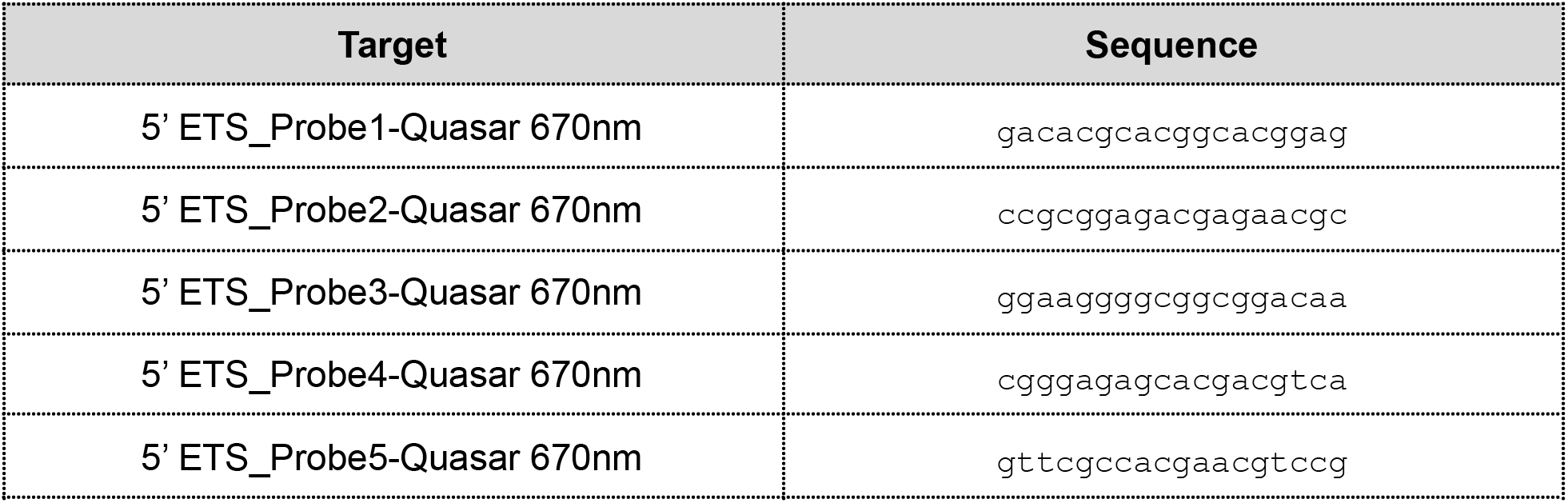
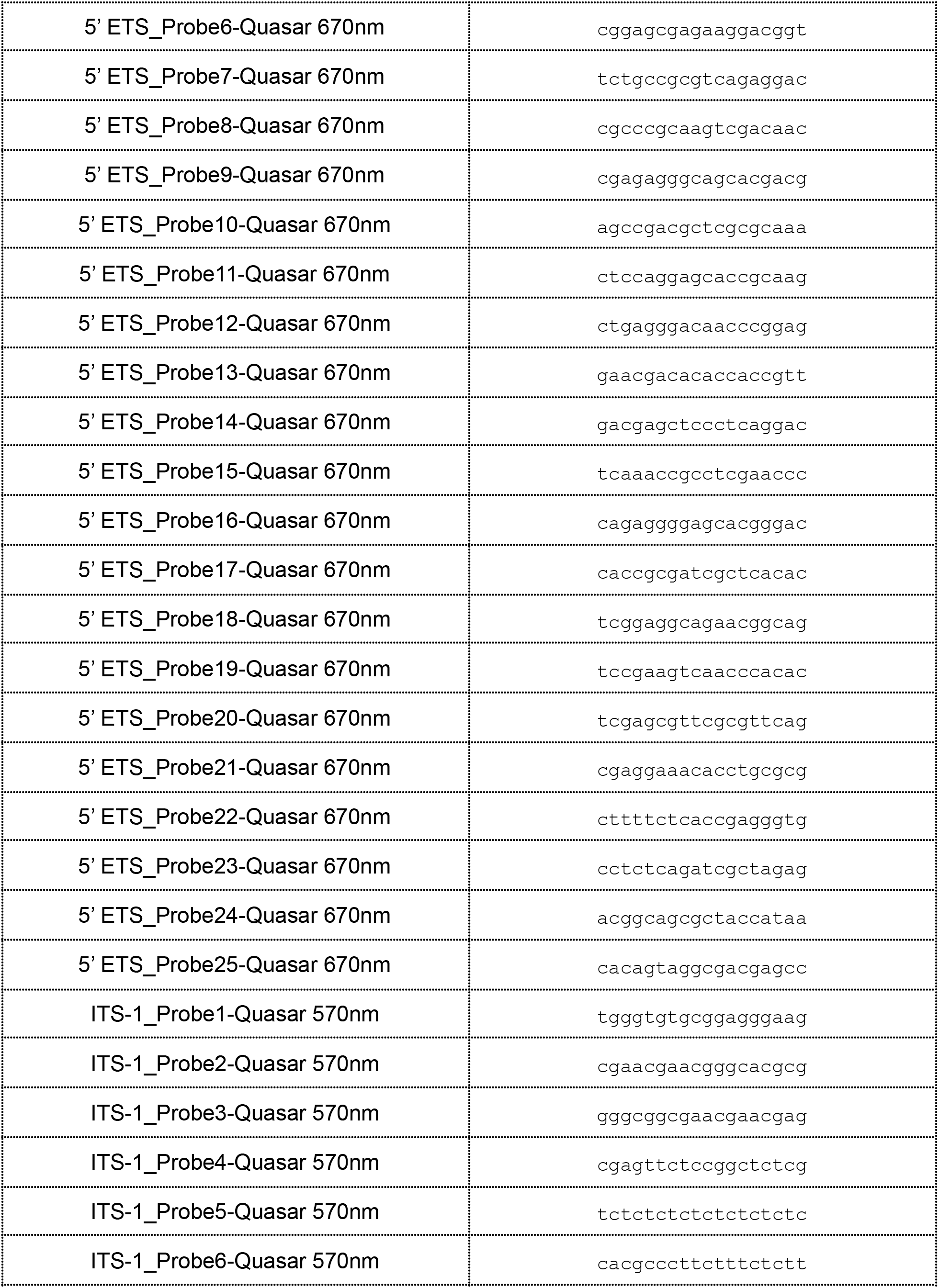
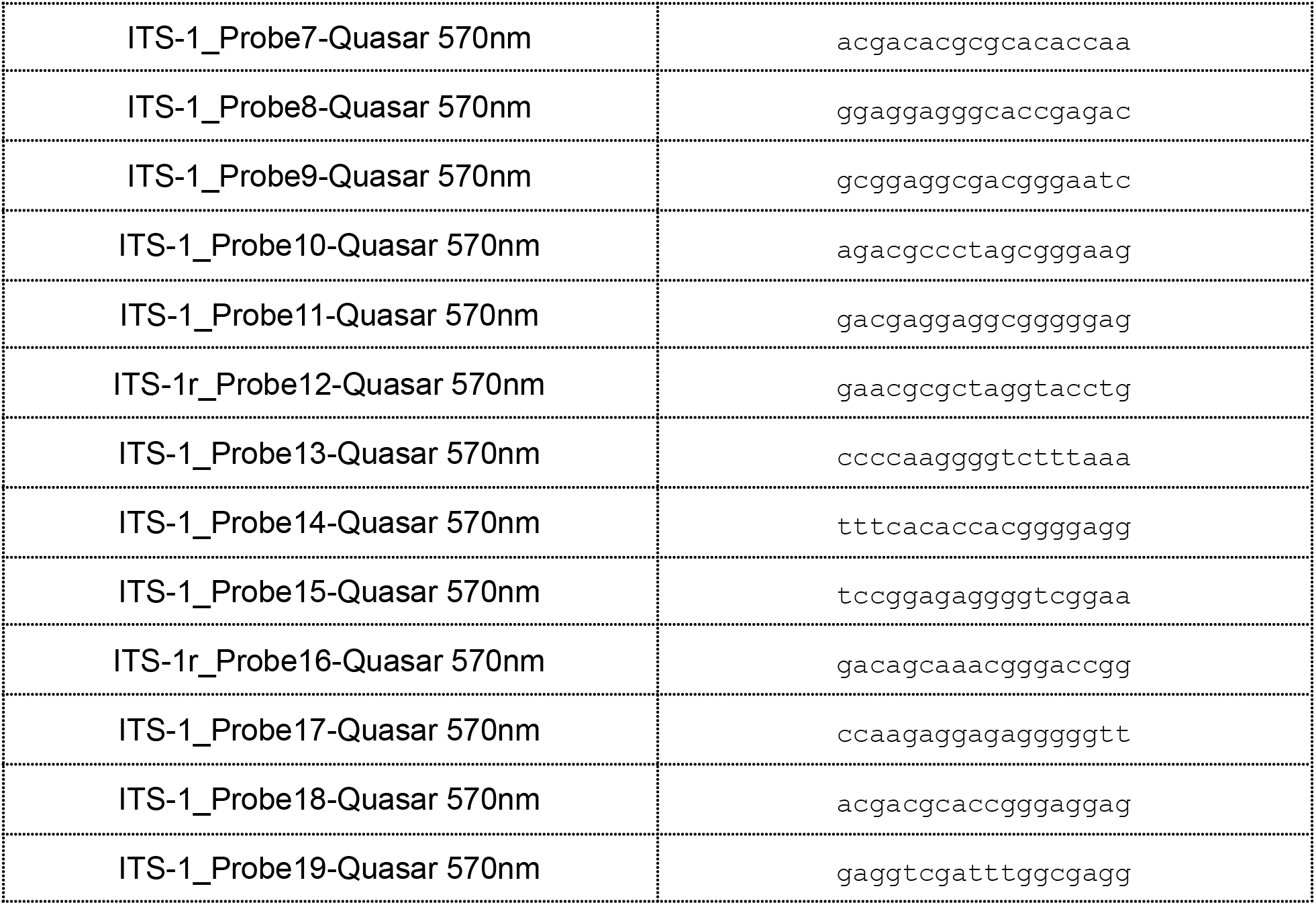
Custom Stellaris RNA-FISH probes.

#### Combined RNA and Protein Imaging

U2OS-Rainbow cells were seeded onto acid-washed #1.5 glass coverslips (at a density of 50,000 cells/coverslip). After 24 hours, the cells were stained with 1 μM HaloTag Oregon Green (Promega) in DMEM for 15 minutes at 37°C and washed with PBS. Cells were then treated for 0, 0.5, 1, 2 and 4 hours with the indicated amounts of drugs in DMEM medium. After fixation with 4% PFA for 15 minutes at room temperature, RNA-FISH staining against the 5’ETS was performed as described above.

Cells were imaged using a Leica SP8 laser-scanning confocal microscope equipped with a 63x (1.4 NA) oil immersion objective. For 3D reconstruction, z-stacks (5.53 μm total in 0.221 μm steps) of selected cells using a 2.5 digital zoom were taken. To remove nuclear background signals, Gaussian filters and image thresholds were used. Images were analyzed and quantified using custom Mathematica scripts.

#### Estimation of NPM1 Surface Tension

UCSF-Rainbow cells were seeded at 15,000 cells/well in an 18-well ibidi glass-bottom μ-slide and allowed to adhere o/n. After treatment with the indicated amounts of drugs for 4 hours, the cells were imaged with an Olympus IX83 epifluorescence microscope equipped with an Orca Fusion scMOS camera, a 100x oil objective (NA 1.45), a X3-ZDC2 TruFocus drift compensator and an Okolab IX3-SVR stage top incubator set to 37°C and 5% CO2. 100 consecutive frames in the FITC channel were taken per field of view at 250 ms intervals and analyzed as previously described (*15*) using custom Mathematica scripts. In short, the surface fluctuation u of individual nucleoli was quantified by averaging the changes in nucleolar contour over both time and polar angle using the Interpolation and Fourier transformation functions of Mathematica. The average surface fluctuation 〈*u*〉 was then used to estimate nucleolar surface tension *γ* based on the relation *γ* = k_B_T/〈*u*^2^〉.

#### Immunofluorescence Stainings and Imaging

U2OS-Rainbow cells were seeded onto acid-washed #1.5 glass coverslips (at a density of 50,000 cells/coverslip). After 24 hours, the cells were stained with 1 μM HaloTag Oregon Green (Promega) in DMEM for 15 minutes at 37°C and washed with PBS. Following treatment with the indicated amounts of drugs, cells were fixed with 4% PFA for 15 minutes at room temperature and permeabilized in blocking buffer (PBS supplemented with 0.5% Triton X-100, 1% donkey serum and 10 mg/ml BSA) for 30 minutes at room temperature. Cells were then incubated with primary antibodies (Table S1; diluted 1:500 in blocking buffer) for 16 hours at 4°C. After washing three-times with PBS containing 0.2% Triton X-100, cells were further incubated with Alexa647-labeled secondary antibodies (Table S1; diluted 1:500 in blocking buffer) for 1 hour at room temperature. Following three washes with PBS containing 0.2% Triton X-100, cells were mounted onto microscopy slides with ProLong Diamond mounting medium containing DAPI (Molecular Probes).

Cells were imaged using a Leica SP8 laser-scanning confocal microscope equipped with a 63x (1.4 NA) oil immersion objective. Images were analyzed and quantified using custom Mathematica scripts.

#### Estimation of ΔG^tr^ and ΔΔG^tr^ for NPM1 and SURF6

15,000 UCSF-Rainbow cells were seeded per well of a 18-well ibidi glass-bottom μ-slide and allowed to adhere o/n. Cells were then treated with the indicated amounts of drugs for 4 hours, fixed with 4% PFA for 15 minutes at room temperature and immunostained for SURF6 as described above. Following washing, cells were kept in PBS and imaged using an Olympus IX83 epifluorescence microscope equipped with an Orca Fusion scMOS camera, a 40x air objective (NA 0.95) and a X3-ZDC2 TruFocus drift compensator. For every well, a 9×9 grid was automatically acquired in the FITC and TRITC channels.

The free energy for the transfer of NPM1 and SURF6 from the dilute phase (nucleoplasm) to the dense phase (nucleoli) was estimated as previously described (*19*). In short, the image segmentation and quantification functions of Mathematica were used to quantify the nucleoplasmic and nucleolar signals, I^dilute^ and I^dense^, for both NPM1 and SURF6. These values were then used to determine the partitioning coefficient *K* = I^dense^/I^dilute^ and the free energy of transfer ΔG^tr^ = - *RT* ln*K*.

#### Cell Synchronization and Live Cell Imaging

1×10^6^ cells were seeded in a 10cm culture dish and incubated for 24 hours. The culture medium was then replaced with fresh DMEM containing 2mM thymidine. After 18 hours, cells were washed twice with PBS, allowed to recover for 6 hours in fresh DMEM, and subjected to a second block with 2mM thymidine for 18 hours.

Synchronized cells were then trypsinized, seeded into 8-well glass-bottom μ-slides (ibidi) at a density of 30,000 cells/well, and allowed to adhere for three hours. Culture medium was then exchanged with L-15 medium (Gibco) supplemented with 10% FBS and the indicated amounts of drugs. Cells were imaged every 15 minutes for a total of 24 hours in a 3×3 grid per well using an Olympus IX83 epifluorescence microscope equipped with an Orca Fusion scMOS camera, a 40x air objective (NA 0.95), a X3-ZDC2 TruFocus drift compensator and an Okolab IX3-SVR stage top incubator set to 37°C. Images were quantified using custom Mathematica scripts.

To determine cell cycle stage, samples were taken immediately after synchronization and at the time point of drug addition, fixed and permeabilized with ice-cold 70% ethanol at 4°C overnight, stained with 1 μg/ml propidium iodine in PBS containing 0.1% Triton X-100 and 10 μg/ml RNase, and analyzed using a Sony SH800 flow cytometer.

#### MTT Assay and Dose-Response Determination

1×10^4^ synchronized cells were seeded per well of two 96-well plates and allowed to adhere for three hours. Cells were then treated with 0, 0.5, 1, 2.5, 5, 7.5, 10, 25, 50, 75, 100, and 200 μM oxaliplatin or cisplatin (four replicates per condition). After 24 and 48 hours, 0.22 μg/μl MTT reagent (3-(4,5-dimethylthiazol-2-yl)-2,5-diphenyltetrazolium bromide) was added per well. After incubation at 37°C and 5% CO_2_ for three hours, the culture medium was removed and cells were solubilized in 50 μL DMSO per well for 5 min at 37°C and 5 min at room temperature. Absorbance (Abs) of the solubilized MTT dye was then quantified at 570 nm and 690 nm using a Biotek Synergy HT microplate reader.

For analysis, the MTT intensity I at concentration c was defined as I(c) = Abs_570nm_(c)-Abs_690nm_(c), and I(c) plotted against c. To determine EC_50_ values, the data was fitted to the equation:

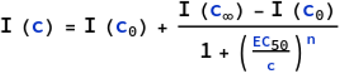

where I(c_0_) is the maximal MTT intensity, I(c_∞_) the minimal MTT intensity and n = 2.

#### Drug Toxicity Rescue Assays

To determine the impact of NPM1 or FBL overexpression on Cis and Ox-induced lethality, HT-1080 Nuc::mKate2 cells were transfected with constructs overexpressing either GFP-NPM1, GFP-FBL, or GFP alone. Briefly, 90,000 cells were seeded into a well of a 6-well plate. Once adherent, media was replaced with antibiotic free growth media for 20 minutes. Then, cells were incubated in a mixture of OptiMEM (ThermoFisher), Lipofectamine LTX (ThermoFisher), and recombinant DNA as per manufacturer protocol. 48 hours after transfection, the cells were lifted and seeded into 384-well plates at a density of 1500 cells/well. The next day, the growth media was changed to fresh media containing cisplatin or oxaliplatin, dosed in a two-fold dilution for a ten-point dose response curve where the highest dose was 100 μM.

Cell viability was then analyzed using the IncuCyte imaging platform, as previously described (*36*). Briefly, the assay measures live cells based on nuclear mKate signals. Images were captured in the red fluorescent mKate channel at 10x magnification every 4 hours for a total of 48 hours post drugging. To count cells, image segmentation was performed using Matlab 9.3.0.713579 (R2017b). Total cells were counted based on threshold for mKate2 levels and transfected cells were counted based on threshold for GFP levels. EC_50_ values were determined as for the MTT assay, except that the concentration-dependent cell number was fitted instead of the MTT intensity.

#### FRAP Analysis

Cells were seeded into 8-well polymer-bottom μ-slides (ibidi) at a density of 30,000 cells/well, allowed to adhere overnight and stained with 1 μM HaloTag Oregon Green (Promega) in DMEM for 15 minutes at 37°C. After washing with PBS, cells were treated with the indicated amounts of drugs in L-15 medium (Gibco) supplemented with 10% FBS. After incubation for 4 hours, live cells were imaged using a Leica SP8 laser-scanning confocal microscope equipped with a 63x (1.3 NA) glycerol immersion objective and a temperature-controlled incubation chamber (Life Imaging Services) set to 37°C. Fluorescence in the Oregon Green and mCherry channels was simultaneously bleached in one nucleolar hemisphere for 10 frames with 488 nm and 561 nm lasers, and the fluorescence recovery in both channels measured every ~1.5 seconds for 60 frames. All half-bleach experiments were performed using the FRAP wizard of the Leica Application Suite X software and analyzed using custom Mathematica scripts.

#### Recombinant Protein Purification

Recombinant FBL and NPM1 were expressed and purified essentially as described previously (*18*). Both FBL and NPM1 were cloned and expressed using the pET system in E. coli BL21(DE3) Rosetta. Transformed cells were grown in LB at 37°C to an OD600 of 0.6 before inducing with 0.5 mM IPTG. Expression continued for up to 4 hours before cells were pelleted and frozen before protein purification.

FBL-expressing cell paste was resuspended in buffer containing 20 mM Tris-HCl, pH 7.5, 500 mM NaCl, 10 mM imidazole, 14 mM β-mercaptoethanol, 10% (vol/vol) glycerol and a protease inhibitor mixture (Roche Diagnostics). FBL was purified first over a Ni-NTA column and by subsequent purification over heparin resin and polished by size exclusion chromatography. FBL was stored at 10 mg/mL in 20 mM Tris, pH 7.4, 1 M NaCl, 1% (vol/vol) glycerol, and 2 mM DTT.

NPM1-expressing cell paste was resuspended and lysed by sonication in buffer containing 20 mM Tris, 300 mM NaCl, 10 mM β-mercaptoethanol, 20U/mL Benzonase (Millipore), and a protease inhibitor mixture (Roche Diagnostics). NPM1 was purified first over a Ni-NTA column subsequently by anion exchange and polished by size exclusion chromatography. NPM1 was stored at 5 mg/mL in 10 mM Tris, 0.15 M NaCl, 2 mM DTT, pH 7.5.

#### In vitro Platination Assays

Single-use aliquots of FBL were thawed on ice and diluted to 1 mg/mL in 20 mM Tris-HCl pH 7.4, 100 mM NaCl, and 100 μg/mL nucleic acid in the presence or absence of platinum compounds. Similarly, NPM1 platination assays were performed by diluting single-use aliquots of NPM1 to 0.5 mg/mL in 10 mM Tris, 0.15 M NaCl, 2 mM DTT, pH 7.5, and 100 μg/mL nucleic acid. Planination of nucleic acid and/or nucleolar proteins was carried out at 25°C for 20 hours in the dark. Total RNA was prepared by phenol-chloroform extraction from HeLa cell pellets. rRNA was purchased from BioWorld. ssDNA source was 20/100 oligo length ladder (IDT), ssRNA source was RiboRuler Low Range RNA standard (Thermo Fisher) and dsDNA ladder was 1kb Plus DNA ladder (NEB).

Protein and nucleic acid platination was assessed by gel shift assays using SDS-PAGE (protein), standard AGE (dsDNA), urea-AGE (ssDNA) and Hepes/triethanolamine-formaldehyde AGE (ssRNA). Proteins were detected by immunoblotting using antibodies against FBL and NPM1 (see Table 1). Nucleic acids were stained with SYBR Green.

## Supplementary Figures

**Figure S1.**
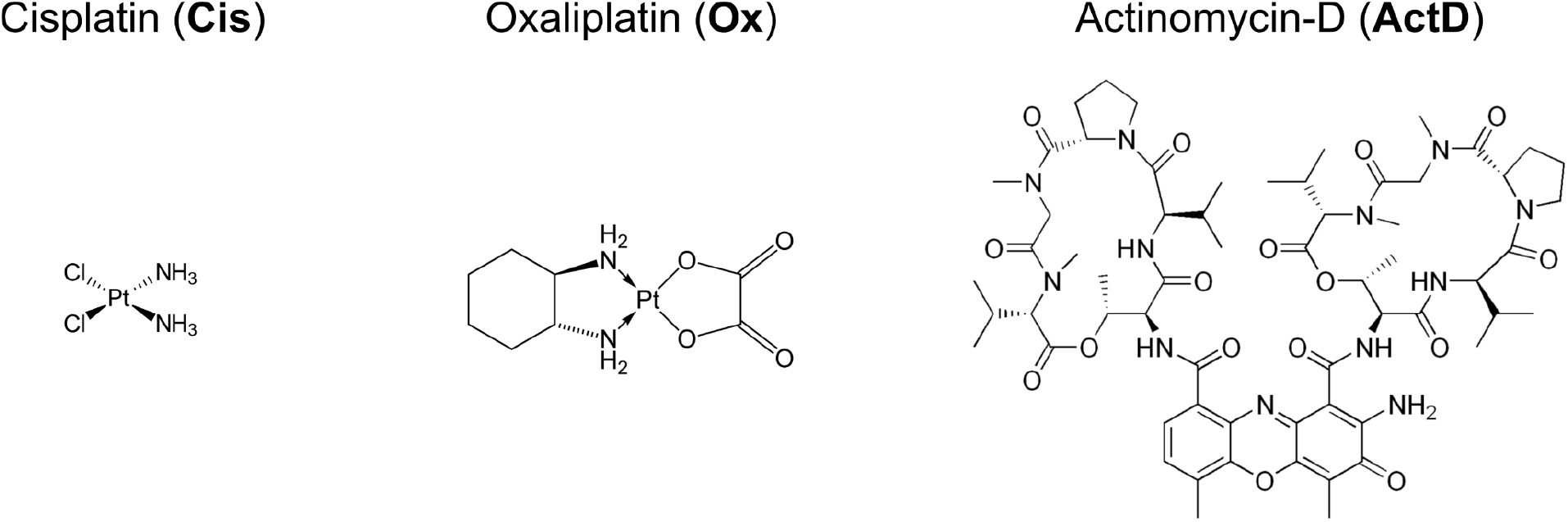
Chemical structures of the small molecule drugs used in this study.

**Figure S2.**
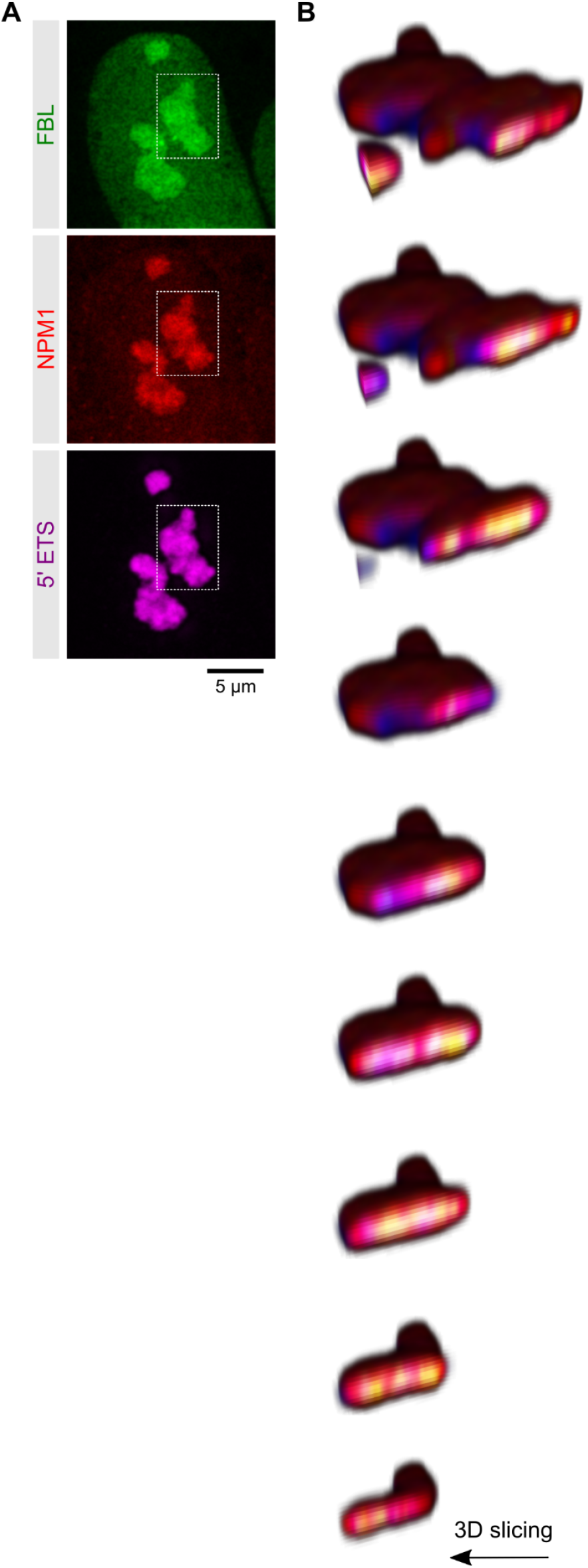
Nucleolar sub-organization in U2OS Rainbow cells. (**A**) Confocal images of the nuclear plane showing the two-dimensional nucleolar sub-organization of NPM1 (red), FBL (green) and the 5’ETS signal (blue). Proteins were endogenously tagged with fluorescent markers and the 5’ETS was visualized using FISH. (**B**) Slicing through the three-dimensional reconstruction of the nucleolus outlined in (A) to illustrate the nucleolar sub-organization of the NPM1 (red), FBL (green) and 5’ETS (blue) layers.

**Figure S3.**
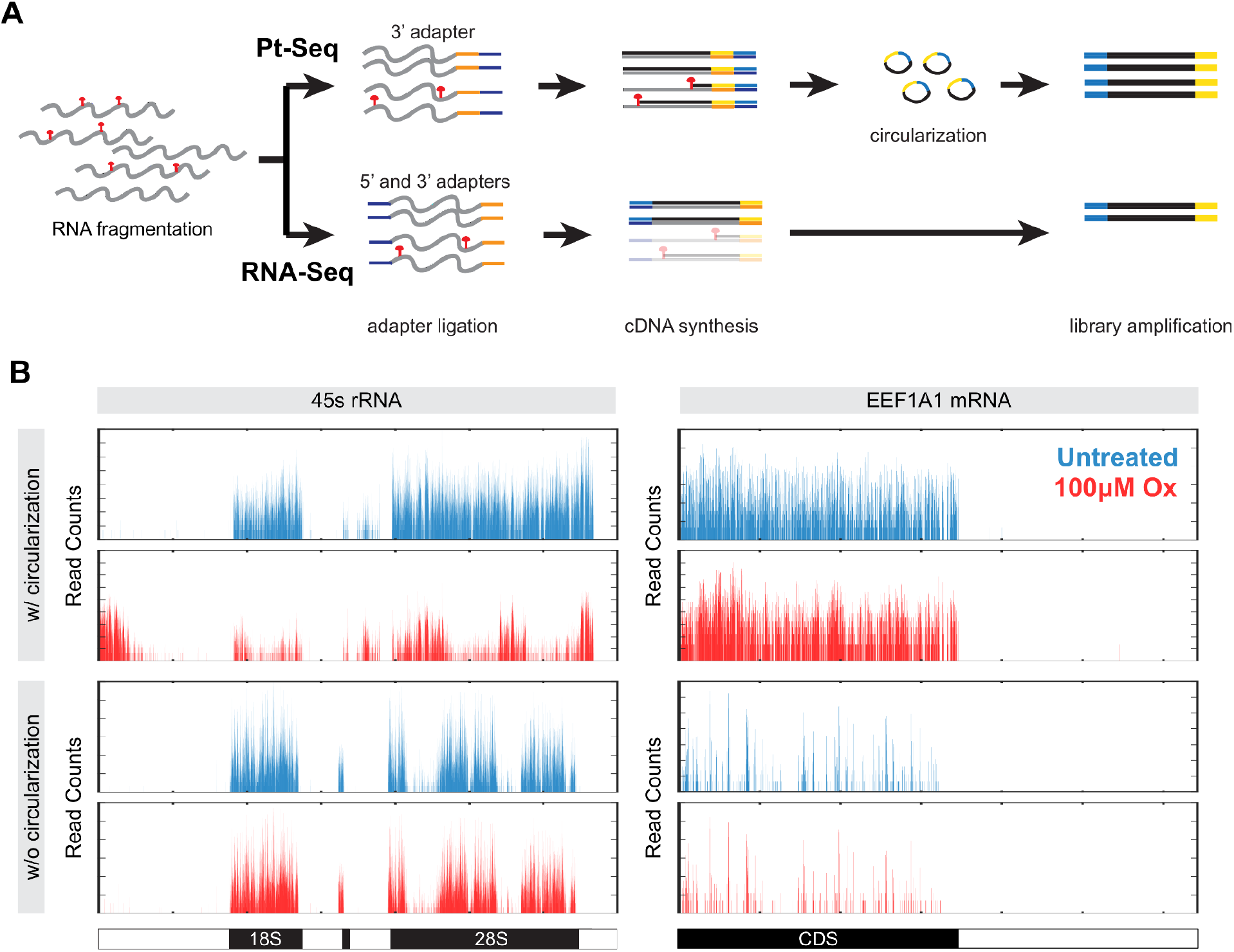
Identification of Pt-modified RNA transcripts in oxaliplatin-treated cells. (**A**) Schematic comparing the workflow of conventional RNA-seq and RNA-seq with circularization, as in Pt-Seq protocols. Due to the circularization step, platinated transcripts can be sequenced and counted. (**B**) Reads from RNA-seq with and without circularization mapped to the 45S rRNA and EEF1A1 mRNA in untreated (blue) and oxaliplatin-treated (red) Huvec cells.

**Figure S4.**
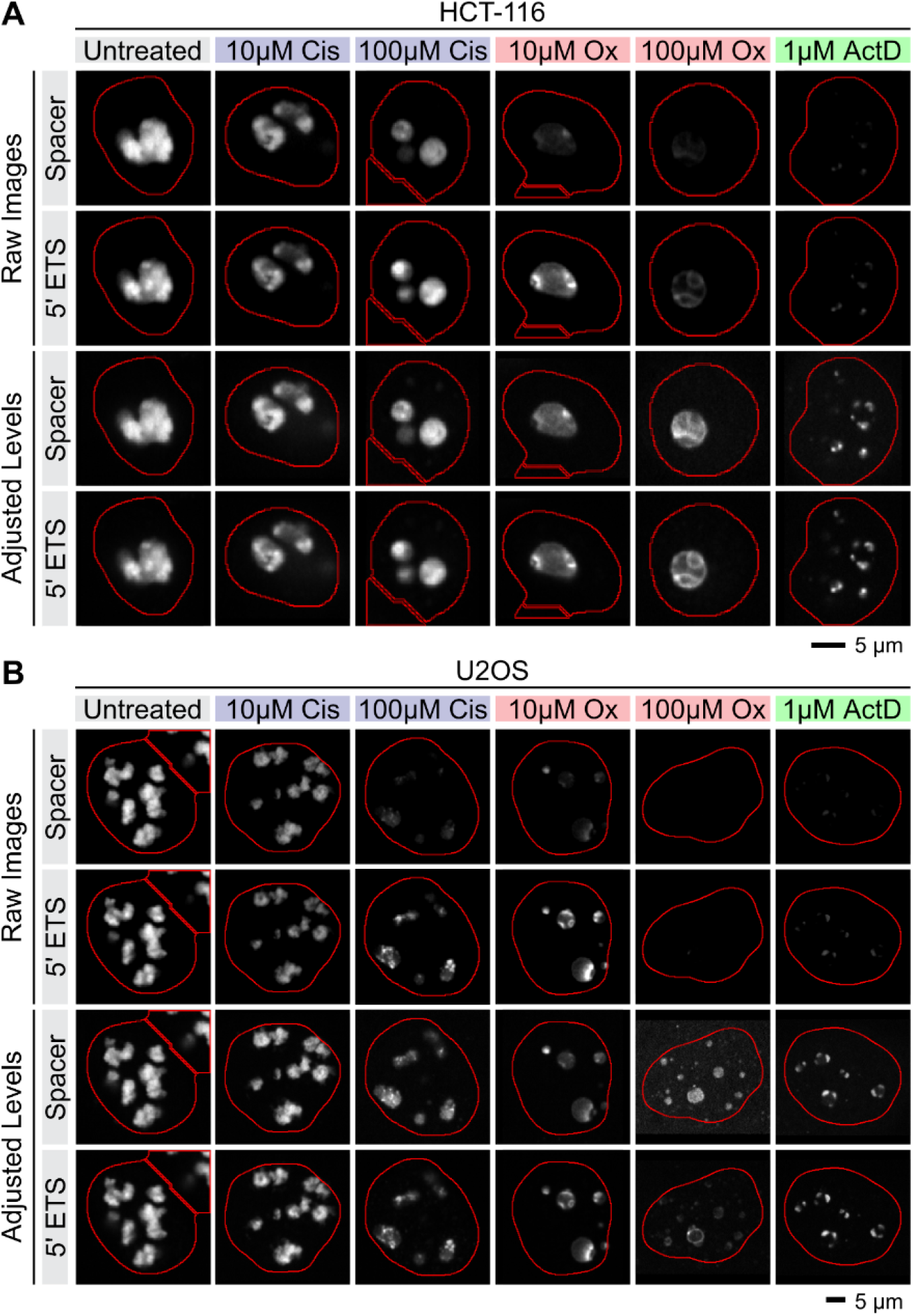
RNA-FISH staining against the 45S rRNA in cancer cell lines. Shown are the individual channels of the raw and adjusted images used for the overlays in Figure 1E. Stainings were performed in HCT-116 (**A**) and U2OS (**B**) cells.

**Figure S5.**
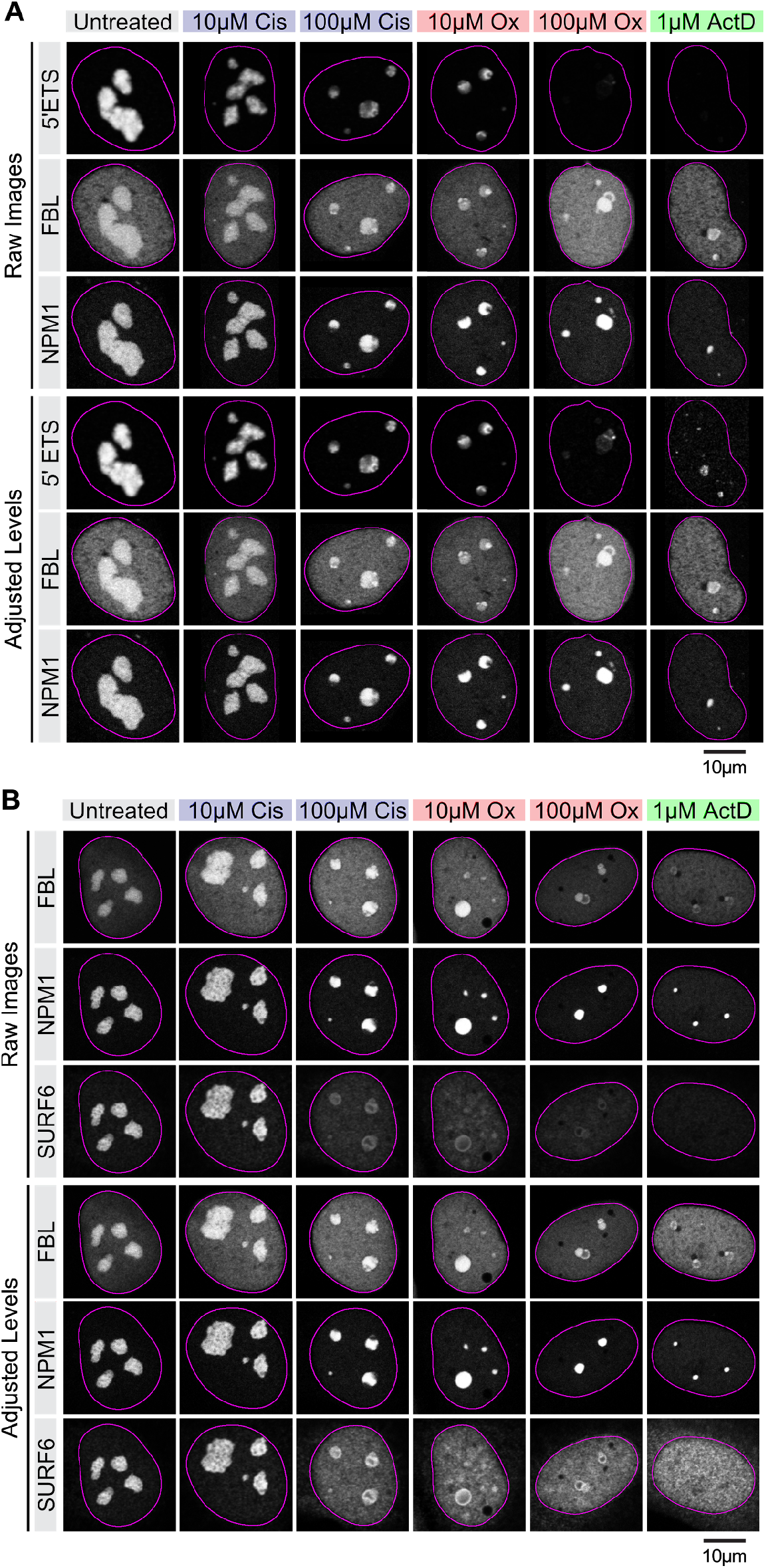
Nucleolar demixing in response to drug treatments. (**A**) Raw and adjusted images used in Figures 2A. (**B**) Raw and adjusted images used in Figure 2D.

**Figure S6.**
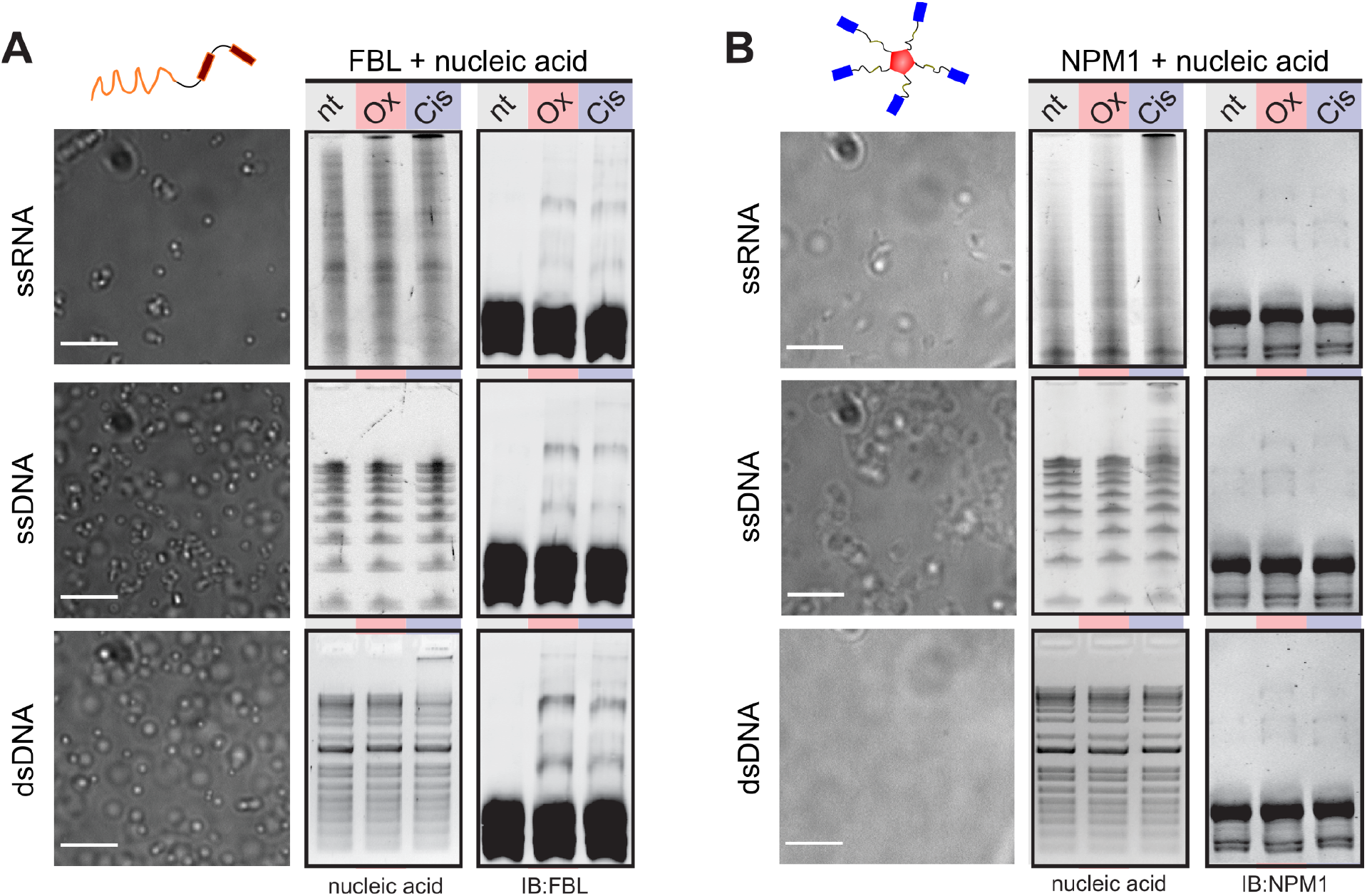
Biochemical analysis of nucleic acid and protein modification by cisplatin and oxaliplatin. A combination of DIC imaging, Hepes/triethanolamine-formaldehyde AGE (ssRNA), urea-AGE (ssDNA), standard AGE (dsDNA), and SDS-PAGE (protein) was used to assess nucleic acid-protein phase separation and platination of nucleic acids and proteins, respectively. Nucleic acids were stained with SYBR Green and proteins detected by immunoblotting. (**A**) Fibrillarin (FBL) readily phase-separates in the presence of ssRNA, ssDNA and dsDNA. Note that FBL is platinated by both oxaliplatin and cisplatin in vitro. Cisplatin also modifies ssRNA and dsDNA. (**B**) Nucleophosmin (NPM1) may phase-separate in the presence of ssRNA and ssDNA. Condensation may enhance the platination of the nucleic acids by cisplatin. Scale bars: 5 μm.

**Figure S7.**
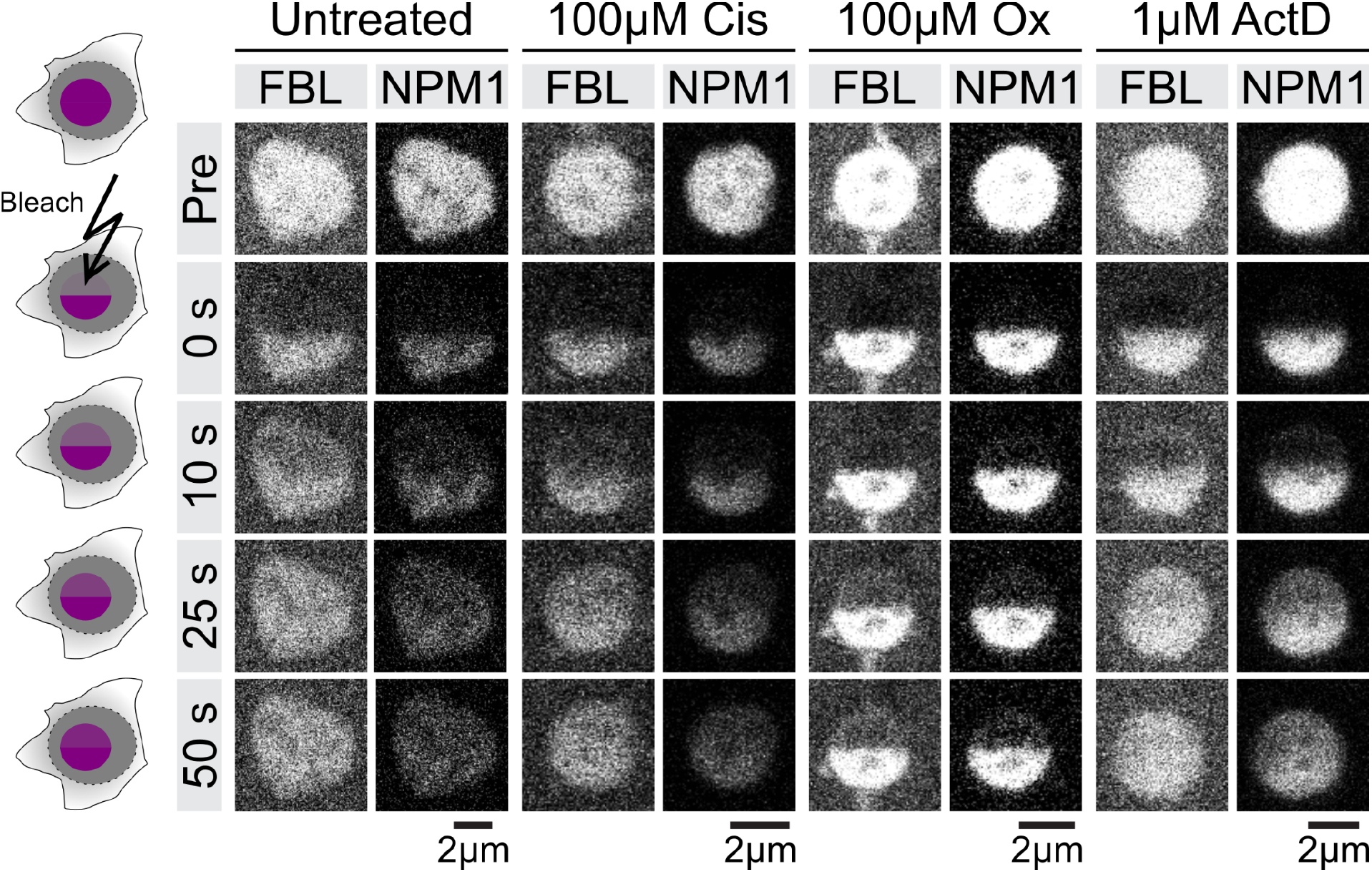
Half-bleach fluorescence recovery after photobleaching analysis to measure nucleolar dynamics. Left: Schematic illustrating the experimental workflow. Right: Raw images used for the overlays in Figure 3E.

**Figure S8.**
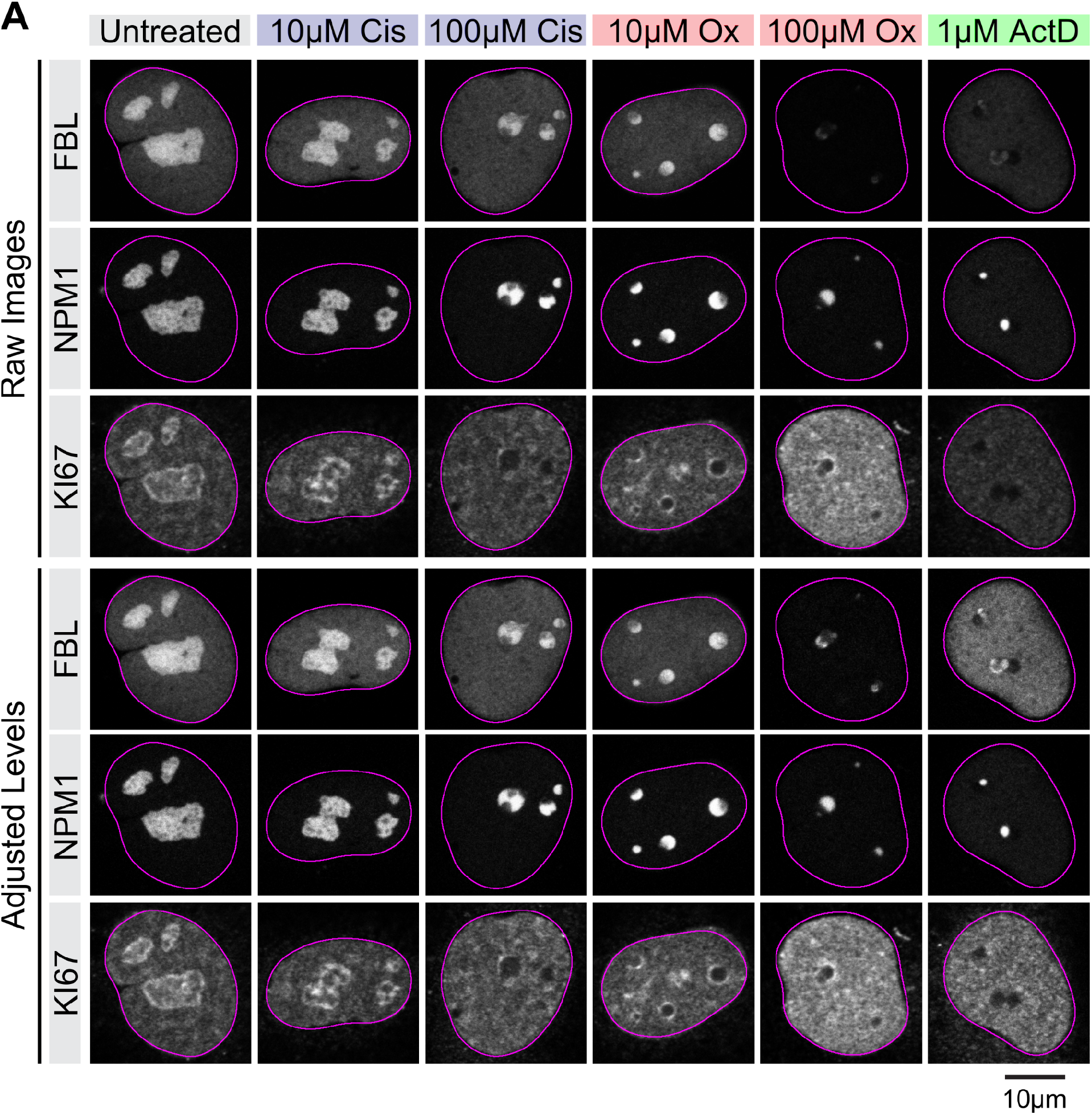
Immunofluorescence staining against KI67 in U2OS Rainbow cells. (**A**) Shown are the raw and adjusted images used in Figure 4A.

**Figure S9.**
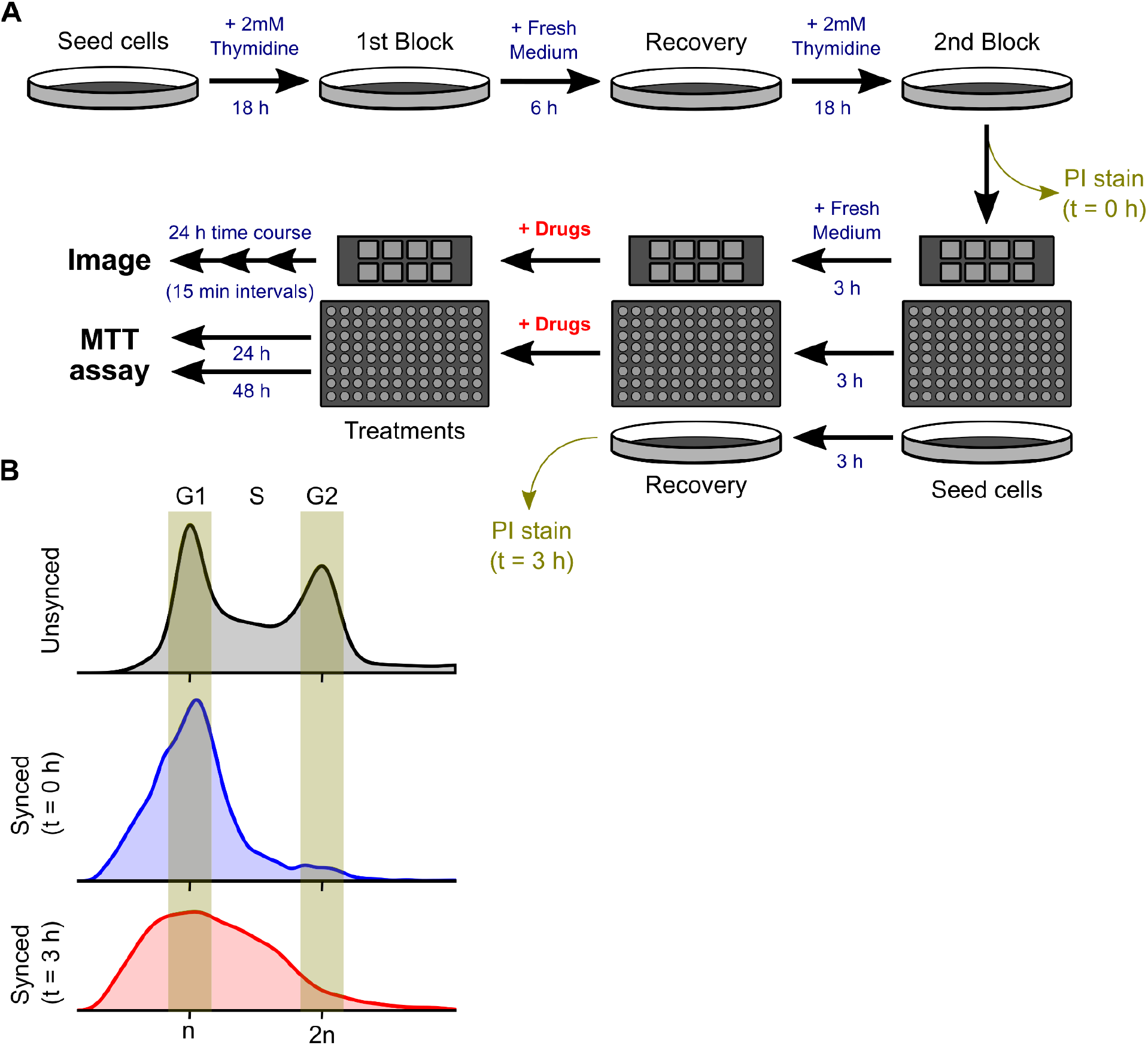
Synchronization of U2OS Rainbow cells. (**A**) Flow chart outlining the synchronization process using the double-thymidine block method. After the second block, cells were seeded for analysis by live cell imaging, MTT assay and flow cytometry. (**B**) Histograms showing the DNA content of unsynchronized and synchronized cells stained with propidium iodide and analyzed with flow cytometry as outlined in (A).

**Figure S10.**
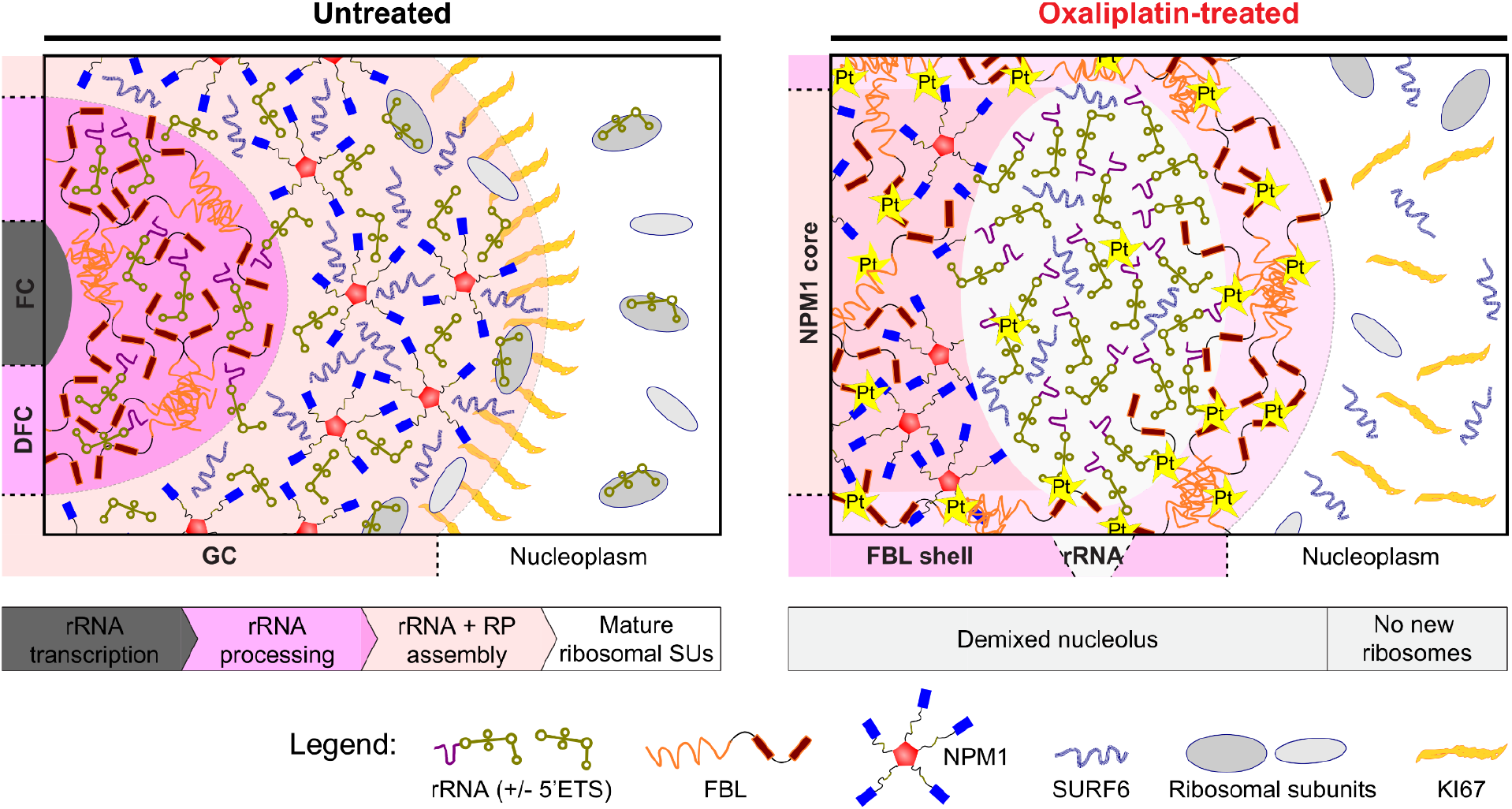
Model of oxaliplatin action. Cartoons illustrate the sub-nucleolar organization of untreated (left) and oxaliplatin-treated (right) nucleoli and the key molecular interactions underlying this organization. Flow charts below the cartoons map the process of vectorial ribosome assembly to the sub-nucleolar layers and how this process is disrupted in oxaliplatin-treated nucleoli.

